# Relating Bone Strain to Local Changes in Radius Microstructure Following 12 Months of Axial Forearm Loading in Women

**DOI:** 10.1101/2020.06.10.144634

**Authors:** Megan E. Mancuso, Karen L. Troy

**Affiliations:** Department of Biomedical Engineering, Worcester Polytechnic Institute, 100 Institute Road, Worcester, MA 01609

## Abstract

Work in animal models suggest that bone structure adapts to local bone strain, but this relationship has not been comprehensively studied in humans. Here, we quantified the influence of strain magnitude and gradient on bone adaptation in the forearm of premenopausal women performing compressive forearm loading (n=11) and non-loading controls (n=10). High resolution peripheral quantitative computed tomography (HRpQCT) scans of the distal radius acquired at baseline and 12 months of a randomized controlled experiment were used to identify local sites of bone formation and resorption. Bone strain was estimated using validated finite element (FE) models. Trabecular strain magnitude and gradient were higher near (within 200 µm) formation versus resorption (p<0.05). Trabecular formation and resorption occurred preferentially near very high (>95th percentile) versus low (<5th percentile) strain magnitude and gradient elements, and very low strain elements were more likely to be near resorption than formation (p<0.05). In the cortical compartment, strain gradient was higher near formation versus resorption (p<0.05), and both formation and resorption occurred preferentially near very high versus low strain gradient elements (p<0.05). At most, 54% of very high and low strain elements were near formation or resorption only, and similar trends were observed in the control and load groups. These findings suggest that strain, likely in combination with other physiological factors, influences adaptation under normal loads and in response to a novel loading intervention, and represents an important step toward defining exercise interventions to maximize bone strength.

## INTRODUCTION

Osteoporotic fractures represent a significant clinical burden, with 1 in 3 women over age fifty experiencing a fragility fracture in their lifetime [1]. Exercise may have the potential to increase bone mass and offset age-related bone loss. Athletes applying high-intensity mechanical loads over extended periods of time have higher bone density than their peers [2,3], and develop site-specific loading adaptations such as increased cortical thickness in the dominant arms of tennis [4,5] and baseball [6] players. In normal healthy adults, clinical trials have shown that high-impact and resistive exercises consistently elicit a 1-3% increase in bone density at the hip over 6-24 months [7–10]. However, moving towards a more personalized approach that tunes interventions to deliver the optimal “dose” of loading for an individual requires a detailed understanding of the relationship bone tissue loading and changes in bone structure.

Animal models have provided insight into the local relationship between bone loading and adaptation. Early models established a controlling role of mechanical strain magnitude [11,12] and the novelty of strain distribution [13,14] on the amount of new bone formed following a dynamic loading intervention. Focusing on the local relationship between bone strain and adaptation, it has been shown that cortical bone formation is related to local strain magnitude [15] and spatial gradient [16,17]. In murine vertebral loading models focusing on the trabecular bone response, principal stresses, principal strains, strain energy density, and the spatial gradient of strain moderately predict the initiation of trabecular bone formation and resorption [18–21]. It is generally suggested that these tissue-level deformations drive cellular-level mechanical stimuli, such as fluid streaming potentials and membrane shear stresses, which are transduced by osteocytes into biochemical cues regulating osteoblast and osteoclast activity [22,23]. While the mechanisms linking bone strain and cell behavior are likely similar in humans, the strength of the relationship between strain and adaptation may differ due to greater genetic variability and the influence of systemic factors such as hormones, diet, and exercise history. However, due to technical challenges in measuring bone strain and changes in bone microstructure *in vivo* in humans, data addressing this question are extremely limited.

Previously, we established a voluntary forearm loading model [24], in which human participants lean onto their palm to compress a padded load cell that provides feedback to guide load magnitude. We also validated participant-specific finite element (FE) models [25] of the forearm to simulate this loading task, enabling us to characterize the strain environment within the radius bone. These methods can be combined to prospectively assign and measure radius bone strain during the axial loading task. In a previously published pilot study that included twenty-three women, we found that 14-week changes in integral bone density in the distal radius, divided into local regions by quadrant, were positively correlated with continuum FE-estimated energy equivalent strain [26]. While these results provide preliminary support for local, strain-driven adaptation, the continuum FE models do not explicitly consider the effect of trabecular microstructure.

High-Resolution Peripheral Quantitative Computed Tomography (HRpQCT) allows for the *in vivo* imaging of human bone microstructure in ∼1 cm sections of the distal radius [27]. Applying this technology, we validated a multiscale modeling approach [28], which incorporates a micro-FE section based on HRpQCT into continuum forearm FE models. Here, we used this technique and serial HRpQCT imaging to investigate the relationship between tissue-level bone strain and local bone adaptation in the distal radius of healthy, premenopausal women participating in a 12-month, prospective study using our forearm loading model. Our overall hypothesis was that bone formation occurs preferentially in high-strain (magnitude and gradient) regions, while bone resorption occurs preferentially in low-strain regions.

## METHODS

### Participants and Loading Intervention

The data reported here were collected as part of a larger, institutionally approved randomized controlled trial enrolling 102 women [29]. Full enrollment criteria are reported elsewhere [30]. Briefly, women ages 21-40 with healthy BMI, menstrual cycles, serum vitamin D levels, calcium intake, and forearm areal bone density were included. Women were excluded if they had a history of medical conditions or use of medications affecting bone metabolism, a previous injury to the non-dominant arm, or regularly participated in activities applying high-impact loads to the forearm. The current analysis includes a subset of twenty-one participants from the control (non-loading, n=10) and loading (n=11) groups. Control subjects with high-quality HRpQCT (motion ≤ 3) [31] scans available at baseline and 12 months we included. In addition to having good quality scans, we further limited the load group to individuals in the top 50% of participants for achieved loading dose and who were “responders,” meaning they experienced increases in bone density above the least significant change (details below).

The purpose of the parent study was to determine the influence of average strain magnitude and strain rate within the distal radius on changes in average bone structure parameters. Participants were randomized into either a non-intervention control group, or one of several loading groups with a range of target strain magnitudes and loading rates. The applied force required to generate the desired strain magnitude within the distal radius was determined for each individual using subject-specific continuum finite element models. Loading was performed on a custom device with visual LED cues to guide force magnitude and auditory beeps to guide loading rate (Fig 1a). Participants were asked to complete three sessions of axial compressive loading of their non-dominant forearm per week. Achieved loading was monitored by the device, which included a uniaxial load cell (Standard Load Cells; Gujarat, India) and data logger (DATAQ DI-710) to record applied vertical force at 100 Hz. Load cell signals were analyzed in Matlab, and an overall “loading dose” was calculated for each participant as the product of average peak force [N], average loading rate [N/s], and number of loading sessions performed. For the present study, loading participants were ranked by loading dose and only the top 50% were included.

**Fig. 1.**
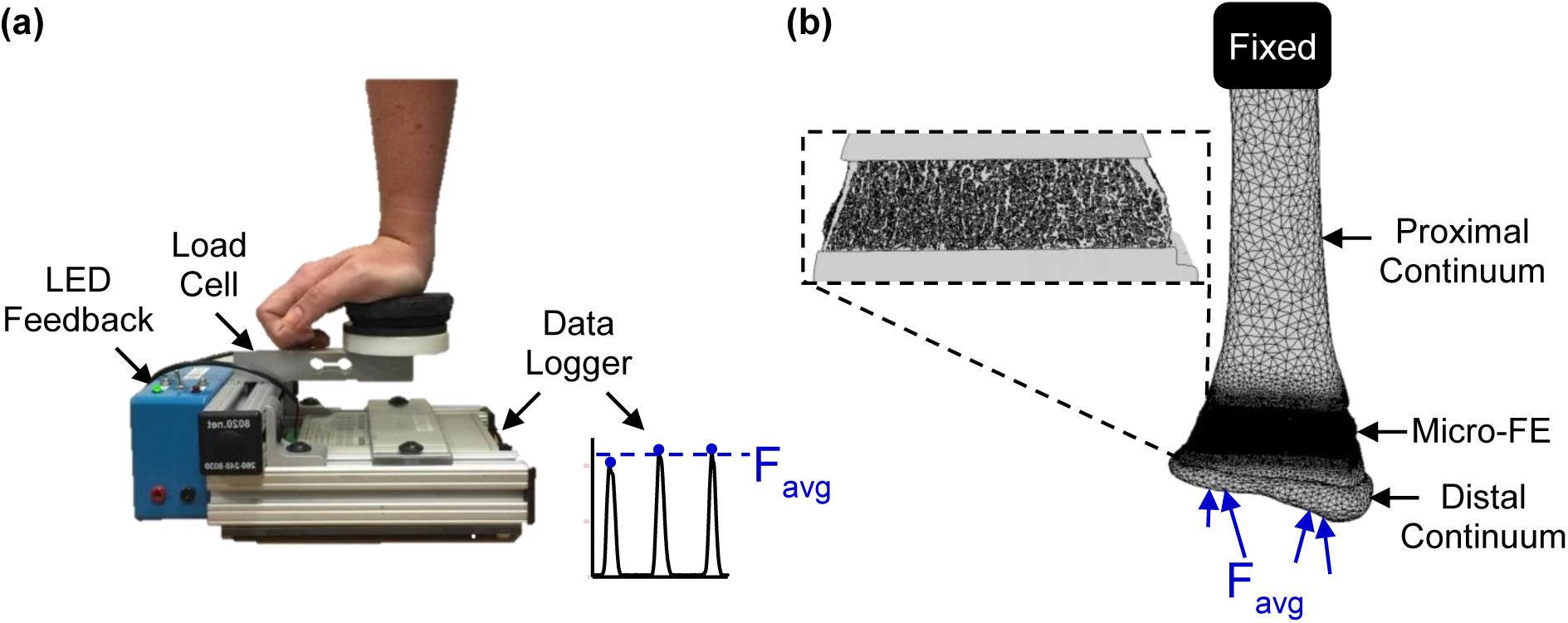
(a) Loading device used to perform forearm loading task. The vertical force was recorded and used to calculate the average applied force, *F*_*avg*_, for each participant in the load group. (b) Multiscale FE models were generated from participant-specific CT scans. For the load group, an axial force equal to the participant-specific average was applied at the distal end. For the control group, the applied force was equal to the overall average across the load group participants.

### Measurement of Bone Adaptation

Local regions of bone formation and resorption within the distal radius were identified from HRpQCT scans (Xtreme CT I, Scanco Medical; Brüttisellen, Switzerland) acquired at baseline and twelve months (isotropic voxel size: 82 m, 0.9 mA, 60 kV). Scans included 110 axial slices covering a 9.02 mm region beginning 9.5 mm proximal to a reference line placed at the distal endplate of the radius. Adaptation sites were identified by aligning, subtracting and thresholding the baseline and follow-up greyscale images (Fig. 2) similar to Christen et al. (2014) [32,33]. Three-dimensional rigid registration (Image Processing Language, V5.16, Scanco Medical AG, Bruttisellen, Switzerland) was used to calculate the transformation matrix needed to align the follow-up image to the baseline image coordinates. The baseline and transformed follow-up images were cropped to the mutually overlapping region and subtracted to obtain voxel-by-voxel changes in density, where increases indicate bone formation and decreases indicate resorption. To reduce the effect of noise and other short-term precision errors, the density difference map was thresholded to include only clusters of at least five voxels with differences ≥225 mgHA/cm^3^ as adaptation sites [32]. Adaptation sites were separated into the cortical and trabecular compartment by taking the Boolean intersection of formation and resorption site masks with the trabecular and cortical masks generated by the Scanco Standard Analysis program [34]. To capture periosteal bone formation added outside the baseline bone surface, the cortical mask was dilated seven voxels (574 micrometers) in the transverse plane. A sensitivity analysis of the 11 loading group participants showed that further increasing the dilation size changed the amount of labeled adaptation sites by less than 0.5%.

**Fig. 2.**
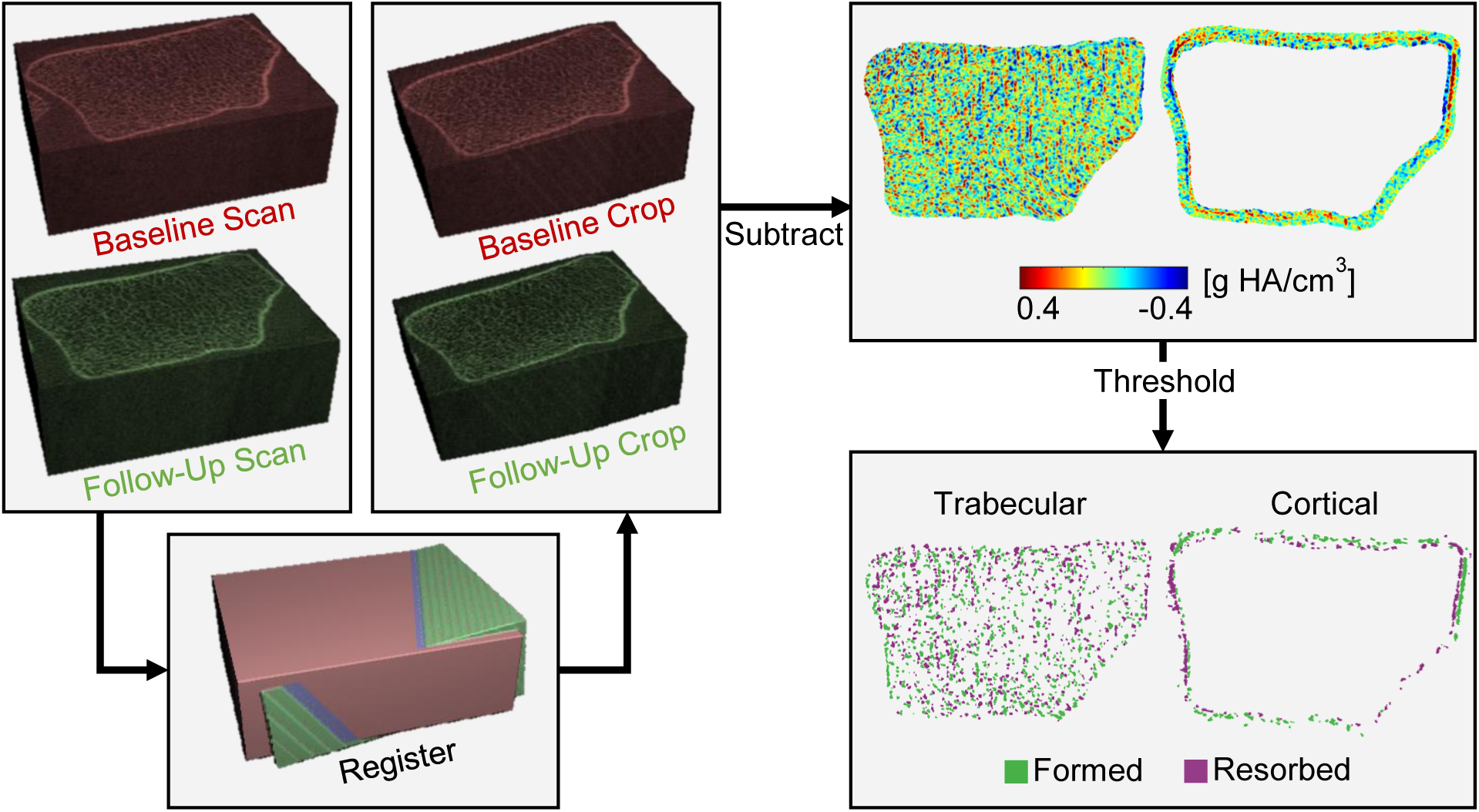
Workflow used to identify local bone formation and resorption sites. Baseline and follow-up HRpQCT greyscale images were aligned and cropped to the overlapping region. Cropped images were subtracted to obtain density difference maps for the trabecular and cortical compartments, which were thresholded to include continuous clusters of at least five voxels with a minimum change of 225 mg HA/cm^3^.

To assess repeatability, this adaptation labeling method was applied to a separate short-term precision data set including eight pairs of repeat distal radius scans acquired within two weeks of each other. The least significant change in average trabecular density for 3D registered scans was calculated as 2.77*CV%_RMS_ [35]. Next, the number of voxels labeled as formation and resorption as a percent of baseline bone volume were then determined for the cortical and trabecular compartments. As no real measurable bone changes are expected within two weeks, these values reflect the amount of erroneously labeled adaptation caused by short-term precision errors. Finally, in our data set, to select “responders,” who we were confident experienced real bone changes, loading participants were included only if they experienced increases greater than this value (1.73%).

### Estimating Local Bone Strain

Bone strain magnitude and gradient were calculated using validated [28], participant-specific multiscale FE models. These models include the distal 10 cm of the radius, from the wrist joint articular surface to the mid-diaphysis. The radius is divided into two continuum sections (distal, proximal), which flank a micro-FE mesh at the HRpQCT distal radius scanned region (Fig 1a). Continuum mesh regions were generated from baseline clinical resolution CT scans (GE Brightspeed, GE Medical, Milwaukee, WI, in-plane resolution 234 µm, slice thickness 625 µm) of the non-dominant forearm. Three dimensional masks of the scaphoid, lunate, and distal 10 cm of the radius were segmented from the image using a 0.175 g/cm^3^ density threshold [25]. For the radius, the segmented baseline HRpQCT mask was registered to the clinical resolution mask, and regions of the clinical resolution mask outside the HRpQCT region were converted into ten-node tetrahedral FE meshes with a nominal edge length of 3 mm. Continuum radius bone elements were assigned heterogeneous linear elastic material properties (E=1.95 MPa to 35 GPa, v=0.4) based on and established density-elasticity relationship using apparent density [36]. For the micro-FE section, the HRpQCT mask was converted to voxel-based hexahedral elements with an 82 micron edge length and homogenous linear elastic material properties (E=15 GPa, v=0.4). To accurately model radio-carpal contact, a 2 mm thick cartilage surface was generated by dilating the distal radius surface, and modeled with 2 mm ten-node tetrahedrons with hyperelastic neoHookean material properties (E=10 MPa, v=0.45) [37]. The scaphoid and lunate were modeled as rigid non-deformable solids, with ten-none tetrahedral meshes with a nominal edge length of 3 mm.

One cycle of the forearm loading task was simulated in Abaqus CAE (v2016, Simulia, Dassault Systèmes, Vélizy-Villacoublay, France). To reduce computational time, the continuum-only model was used to model full contact at the wrist, which included the scaphoid and lunate carpal bones. Ramped, quasistatic loading was applied through the centroids of each carpal toward the fixed proximal radius, such that the resultant force was axial and equal in magnitude to the participant’s average achieved peak force measured by the loading device. The resulting normal and shear contact forces at radius cartilage nodes were exported from continuum simulations and applied directly at matching nodes in the multiscale model of the radius only. For participants assigned to the control group (who did not apply voluntary loads), a simulation was run with applied force set to the average applied force across all loading participants (324 N). The purpose of the control simulations was to determine an “average” mechanical strain state, to provide a null comparison against the loading group. Multiscale models contained 3,062,520±557,518 nodes and 9,187,560±1,672,555 degrees of freedom (mean ± SD), and simulations were solved on a UNIX server with 54 processors (2.1-3.2 GHz) and 200 GB RAM in 3.4±1.9 hours.

Strain magnitude and gradient were calculated for each element in the distal radius micro-FE region. Principal stresses and strains at element centroids were exported from Abaqus used to calculate energy equivalent strain as

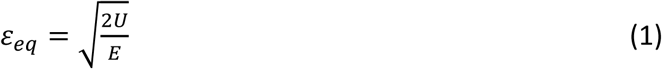

Where *E* is elastic modulus and *U* is strain energy density calculated as

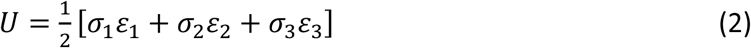

with σ_n_ and ε_n_ being the principal stress and strain components, respectively. Bone strain gradient was calculated as the norm of the gradient of energy equivalent strain in the x, y, and z directions. Gradient in each direction was calculated similar to [38] using the central difference formulation, with simple forward and backward differences calculated for surface elements. For example, gradient in the *x* direction for voxel *i* is calculated as

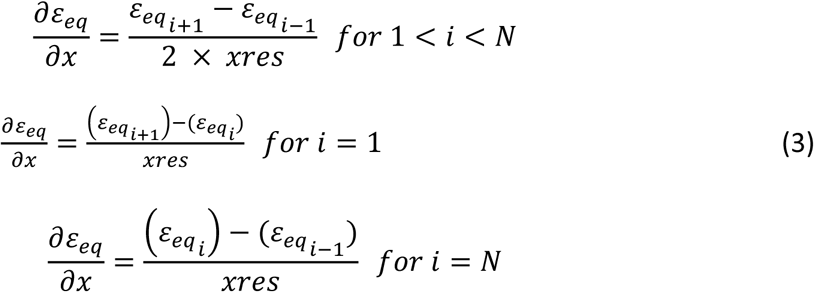

Where *xres* is the element side length (82 microns) and *N* is the number of continuous voxels in the *x*-direction between two surfaces (i.e. of cortical shell or individual trabeculae). The norm of the spatial gradient was calculated for each element as

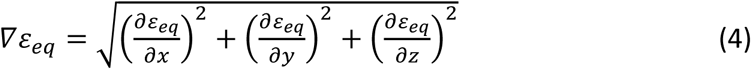

To allow strain and adaptation to be compared spatially, formation and resorption mask coordinates were registered to the multiscale FE coordinates using mutual information 3D registration in Matlab (Mathworks, Natick, MA). A precision analysis demonstrated rotation errors of 0.47 ±0.38°, 0.46 ±0.41°, 0.32 ±0.24° in the x,y,z directions for this method.

### Relating Bone Strain and Adaptation

The hypothesis that bone adaptation is influenced by local tissue strains was tested using four approaches (Table 1). First, we compared strain near sites of formation versus resorption. Second, we compared the percent of bone formation and resorption sites occurring near high versus low strain regions. Third, we compared the percent of high and low strain elements occurring near formation versus resorption. Finally, we characterized the distribution of adaptation and strain within the cortical bone compartment across sixteen angular sectors. All analyses included both the load and control groups.

**Table 1.**
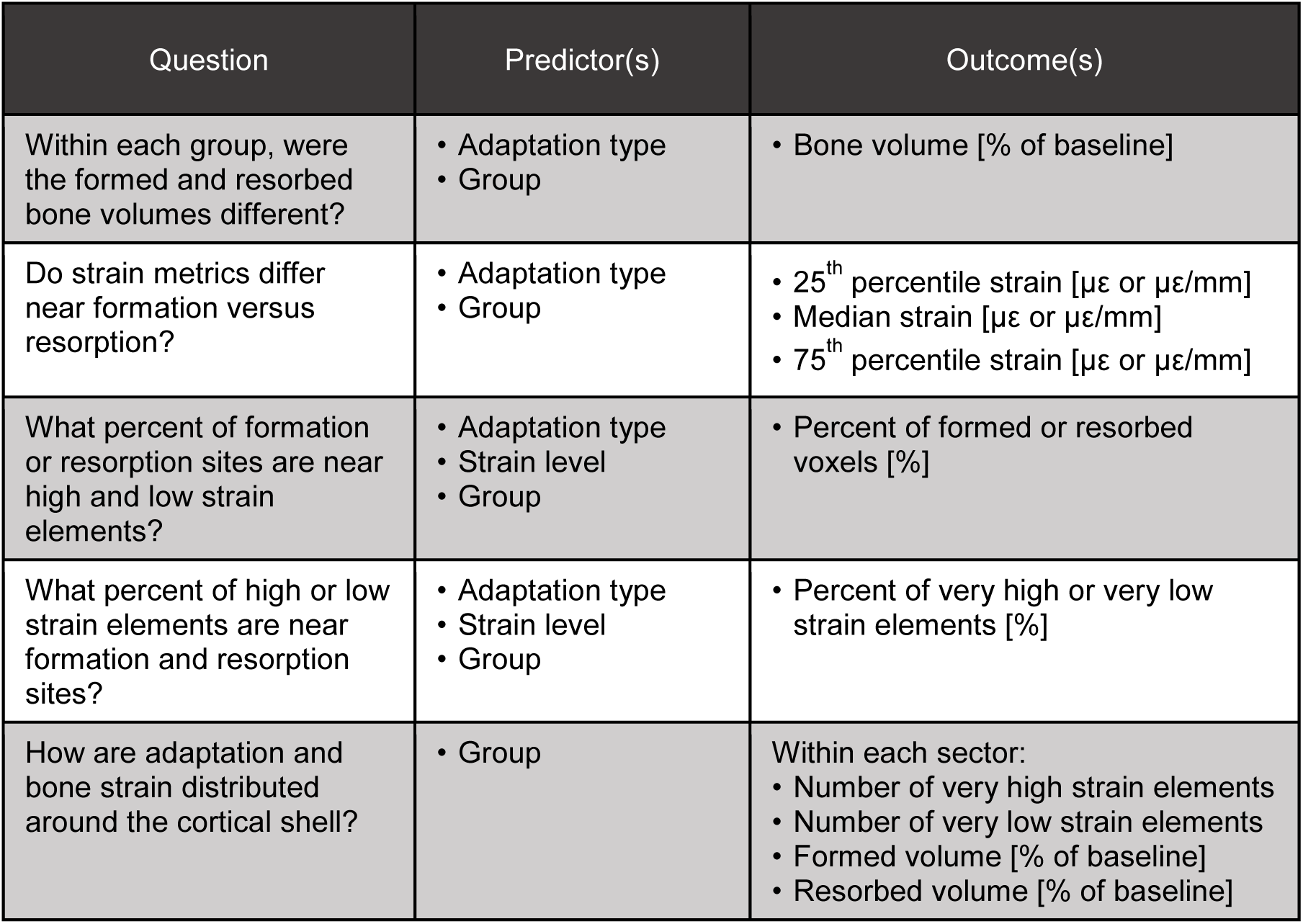
Summary of analyses performed to characterize the amount of bone formation and resorption in each group and spatially relate bone adaptation (formation and resorption) to FE-estimated strain. Each predictor had two levels: formation or resorption for adaptation type, control or load for group, and very high or very low for strain level. For each outcome, separate models were fit for the trabecular and cortical compartments, as well as for strain magnitude and strain gradient for analyses considering strain.

| Question | Predictor(s) | Outcome(s) |
| --- | --- | --- |
| Within each group, were the formed and resorbed bone volumes different? | • Adaptation type<br>• Group | • Bone volume [% of baseline] |
| Do strain metrics differ near formation versus resorption? | • Adaptation type<br>• Group | • 25<sup>th</sup> percentile strain [$\mu\epsilon$ or $\mu\epsilon/\text{mm}$]<br>• Median strain [$\mu\epsilon$ or $\mu\epsilon/\text{mm}$]<br>• 75<sup>th</sup> percentile strain [$\mu\epsilon$ or $\mu\epsilon/\text{mm}$] |
| What percent of formation or resorption sites are near high and low strain elements? | • Adaptation type<br>• Strain level<br>• Group | • Percent of formed or resorbed voxels [%] |
| What percent of high or low strain elements are near formation and resorption sites? | • Adaptation type<br>• Strain level<br>• Group | • Percent of very high or very low strain elements [%] |
| How are adaptation and bone strain distributed around the cortical shell? | • Group | Within each sector:<br>• Number of very high strain elements<br>• Number of very low strain elements<br>• Formed volume [% of baseline]<br>• Resorbed volume [% of baseline] |

For each formation and resorption site, the average strain magnitude and gradient was calculated as the average value for all FE elements within 200 µm, corresponding to 23.8 ±10.3 and 40.0 ±8.3 elements for formation and resorption sites, respectively. Two hundred micrometers was selected as the distance within which osteocyte sense local strains. This falls within the range of previous studies [32,39–41] and is equal to 2.44 element edge lengths. For formation and resorption sites more than 200 µm from a mesh element, the value for the nearest element was used. The median, 25^th^ percentile, and 75^th^ percentile values for average strain magnitude and gradient near formation and resorption in both the trabecular and cortical compartments were determined.

To assess the spatial relationship between adaptation and extreme strain, very high and low strain elements for strain magnitude and gradient were defined using the 9^th^ and 95^th^ percentile values for each metric (Fig. 3). Very high and low strain element sets were defined separately for the trabecular and cortical compartments. The percent of formation sites near very high strain was calculated as the number of formation sites with a high strain element within 200 microns, divided by the total number of formation sites. Conversely, the percent of very high strain elements near formation was calculated as the number of very high strain elements with a formation site within 200 microns, divided by the total number of very high strain elements. Similar calculations were performed for resorption sites and very low strain elements within each bone compartment.

**Fig. 3.**
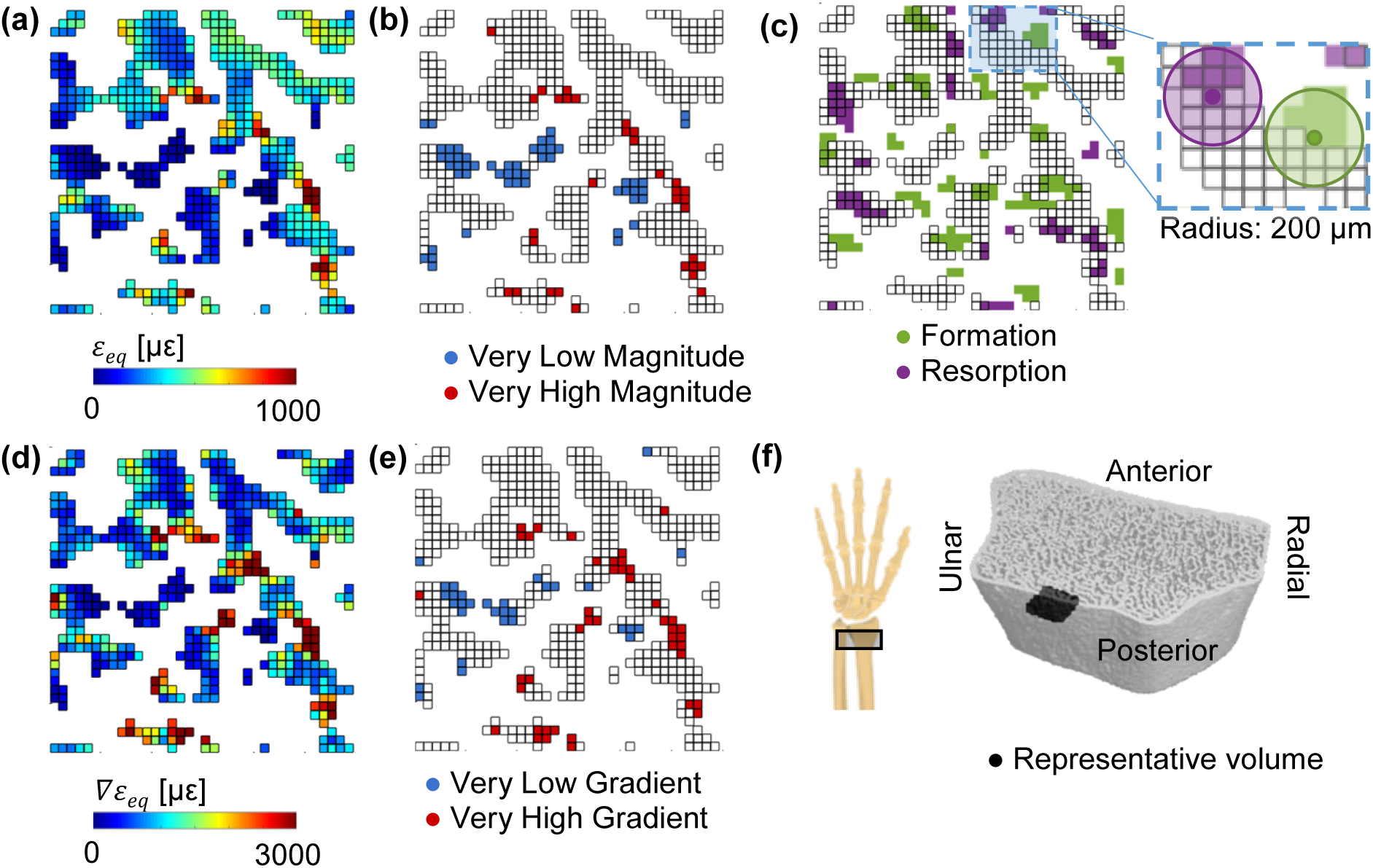
Energy equivalent strain (a) magnitude, *ε*_*eq*_, and (d) gradient, *▽ε*_*eq*_, used to define very low and very high (b) magnitude and (e) gradient elements based on the 5th and 95th percentile values within the trabecular compartment. (c) Formation and resorption sites, with edges indicating elements present in the FE mesh based on the baseline scan. Inset shows 200 µm regions defining which FE elements are near formation and resorption sites. (f) Reconstructed HRpQCT mask of distal radius, indicating the position of the representative 3×3×0.2 mm trabecular volume in black.

To explore the distribution of cortical bone adaptation and strain, we divided the cortical compartment into sixteen equal angle sectors and determined the number of formation versus resorption and high versus low strain elements within each sector (Fig. 4).

**Fig. 4.**
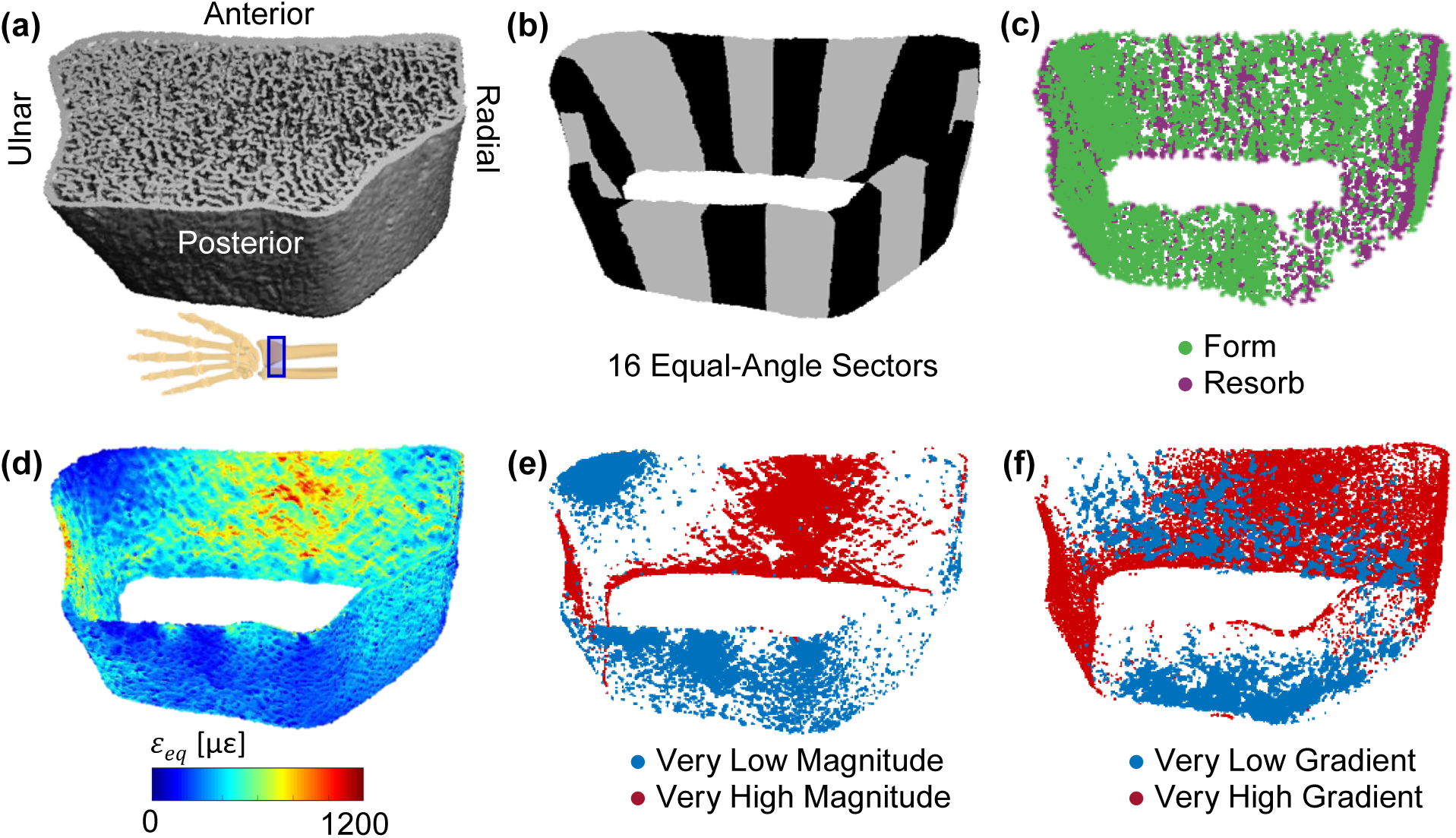
(a) Reconstructed HRpQCT mask of distal radius. (b) The cortical compartment was divided into sixteen equal-angle sectors defined relative to the radius centroid. (c) Cortical formation and resorption sites. (d) Energy equivalent strain within the cortical compartment, used to define very low and high strain (e) magnitude and (f) gradient elements based on 5^th^ and 95^th^ percentile values within the cortical bone compartment.

### Statistics

To characterize adaptation in each group (loading versus control), the volume of formed and resorbed bone within each compartment was compared between groups using a mixed effects linear model. When significant interactions between group and adaptation type were found, formed and resorbed volumes were compared within each group separately.

Strain parameters were compared between adaptation type (formation versus resorption) and group (load versus control) using a mixed effects linear model. Separate models were fit for median, 25^th^ percentile, and 75^th^ percentile strain magnitude and strain gradient within the trabecular and cortical compartments. When significant adaptation by group interactions were found, strain metrics near formation and resorption were compared separately for each group.

The percent of adaptation sites near extreme strain elements was compared using a mixed effects model with adaptation type (formation versus resorption) and strain level (very high versus very low) as repeated measures and group as a between-subjects factor. When significant interactions involving strain level and adaptation type were found, strain level was compared separately within formation and resorption. Similarly, the percent of extreme strain elements near adaptation sites were compared with adaptation type, strain level, and group as factors. When significant interactions involving strain level and adaptation type were found, adaptation type was compared separately within high and low strain elements. Separate models were fit for strain magnitude and gradient within each of the trabecular and cortical bone compartments.

To verify that cortical strains were similarly distributed for load and control groups, independent samples t-tests compared the number of low and high strain elements between groups within each sector. To determine where loading may promote bone formation or prevent resorption within the cortical surface, the number of formation and resorption sites, as a percent of baseline cortical bone volume, was compared between groups within each sector using independent samples t-tests. All statistics were performed in SPSS v25.0, and p<0.05 was used to define statistical significance. Unless otherwise stated, data are expressed as mean±standard deviation.

## RESULTS

### Participants

The participants included in this analysis were 28.7±4.9 years old. On average, the load group participants performed 139±86 sessions of loading over 12 months, applying an average peak force of 324.2±40.7 N.

### Characterization of Local Adaptation

Within the load group, significantly more trabecular bone was formed than resorbed (Fig. 5), consistent with the selection criteria that limited the load group to “responders” with gains in bone density. As a percent of baseline bone volume, 4.1±2.0% more bone was formed than resorbed. Within the control group, formed and resorbed bone volumes were similar, with 1.0±5.5% more bone resorbed than formed.

**Fig. 5.**
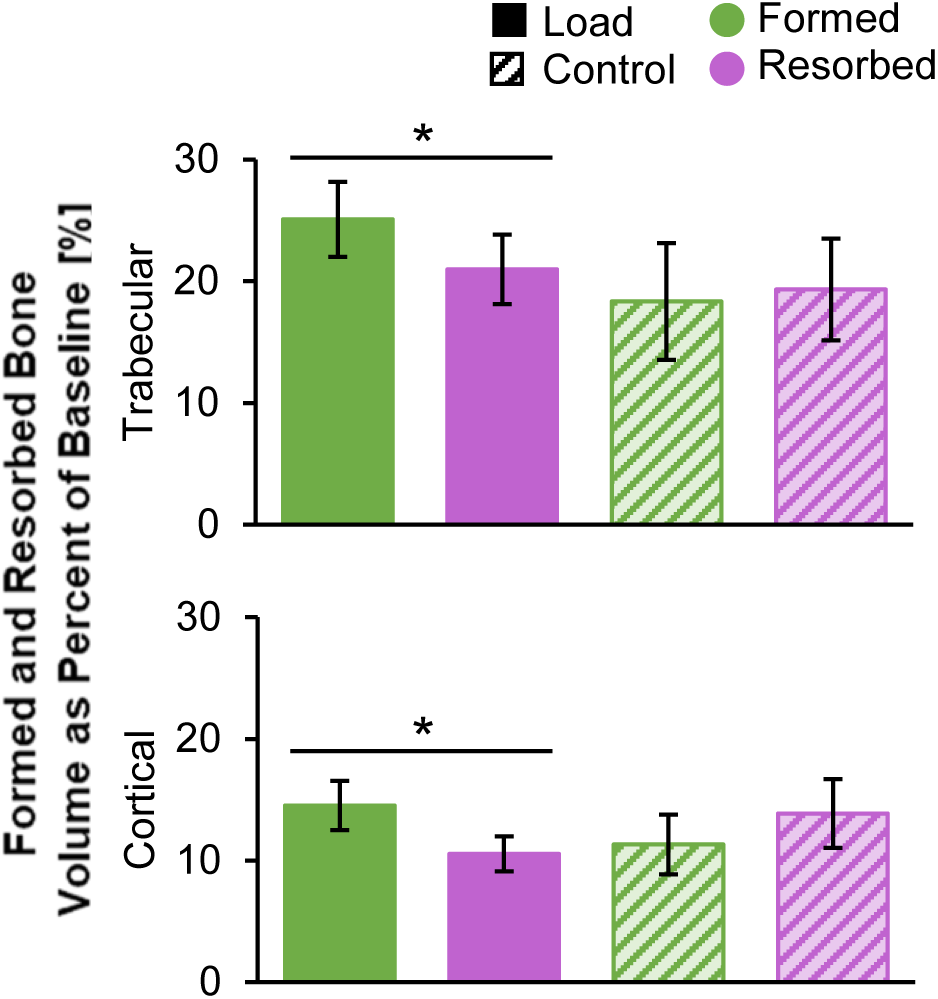
Formed and resorbed bone volume, presented as a percent of baseline bone volume, within the trabecular (top) and cortical (bottom) compartments for the load (n=11) and control (n=10) groups. *Given significant adaptation by group interaction, indicates significant difference between formed and resorbed volume within the load group.

In the cortical bone compartment, the load group experienced significantly more formation than resorption, corresponding to a net increase equivalent to 4.0±4.4% of baseline cortical bone volume. Within the control group, cortical formation and resorption were similar, with 2.5±4.4% more bone resorbed than formed.

Looking at the short-term precision data set, in which no real change occurred, 10.5±4.5% of trabecular bone volume was erroneously labeled as formation and 9.9±4.3% was erroneously labeled as resorption. In the cortical compartment, 4.7±1.9% of baseline bone volume was labeled as formation and 4.9±2.1% was labeled as resorption. The root mean square coefficient of variation for net change for trabecular bone density was 0.62%. Net adaptation within the precision group was not significantly different from zero (p>0.05, one-sample t-test) for both trabecular and cortical compartments, suggesting no systematic bias toward formation versus resorption. The average formed and resorbed volumes in experimental groups were a minimum of 1.7 times those associated with precision error due to partial volume and image registration.

### Do Strain Metrics Differ near Formation versus Resorption?

Trabecular strain magnitude and gradient were higher near formation versus resorption sites for both the load and control groups, except for the 25^th^ percentile of strain magnitude (Fig. 6). While statistically significant, the differences between formation and resorption were relatively small, on the order of 5-10%. This corresponds to an average difference of 11.8±17.2 µε between formation and resorption for median strain magnitude, and 45.7±38.6 µε/mm for median strain gradient (Supplementary Table 1). Thus, the distribution of strain (magnitude and gradient) among formation sites was shifted slightly higher than that of resorption, but the distributions were still mostly overlapping. In the cortical compartment, strain magnitude was similar in formed versus resorbed sites, except at the 25^th^ percentile value, which was higher near resorption. The 25^th^ percentile and median of cortical strain gradient were higher for formation versus resorption in the control group only, with median strain gradient 138.9±98.6 µε/mm higher near formation versus resorption. The 75^th^ percentile of cortical strain gradient was higher for formation versus resorption in both groups. Differences between formation and resorption for cortical strain gradient were relatively high, between 30-40%.

**Fig. 6.**
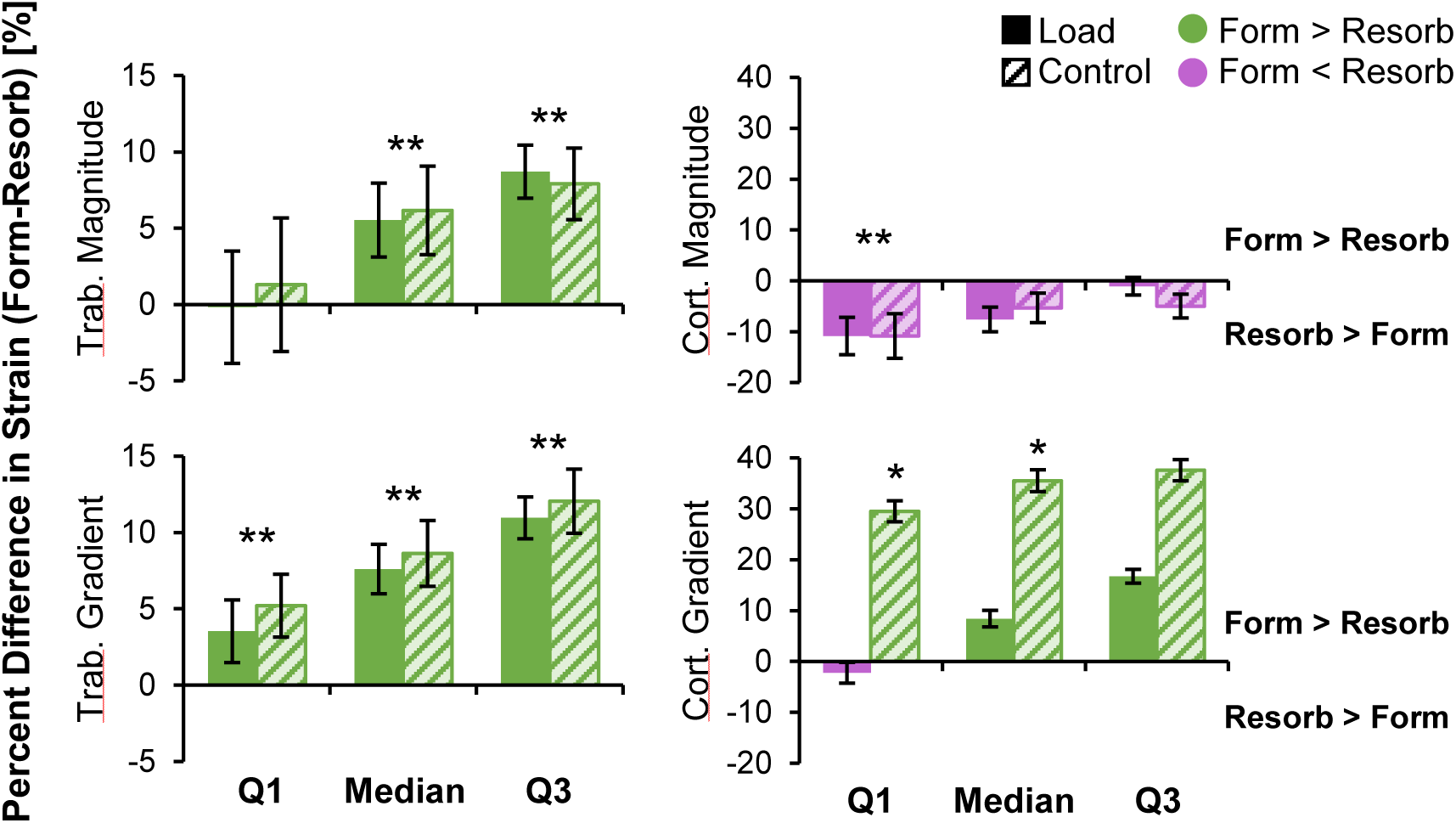
Percent difference in strain metrics between formation and resorption in the trabecular (left) and cortical (right) compartments for the load (n=11) and control (n=10) groups. For each subject, the 25^th^ percentile (Q1), median, and 75^th^ percentile (Q3) of strain magnitude (top) and gradient (bottom) were calculated for formation and resorption. Data presented as group means of within-subject percent difference between formation and resorption (error bars: SEM). Positive differences indicate strain is higher for formation than resorption. *Given significant group by adaptation interaction, indicates significant difference between formation and resorption within the control group only. **Indicates significant difference between formation and resorption for both groups.

### What Percent of Formation or Resorption Sites are Near High and Low Strain Elements?

A greater proportion of trabecular formation and resorption sites were near very high versus very low strain magnitude elements in both the load and control groups (Fig. 7). Similarly, a greater proportion of trabecular formation and resorption sites were near very high versus very low strain gradient elements, particularly for formation sites (p<0.05 for interaction). Cortical bone formation and resorption were both more likely to occur near very high versus very low strain gradient elements. These findings are in line with the hypothesis that high strains lead to microdamage or other biophysical cues that upregulate remodeling, in which both formation and resorption occur. However, while significant differences were observed, over half of formation and resorption sites were near neither very high nor very low strain elements (Supplementary Fig. 1). In the trabecular compartment, a greater proportion of resorption sites were near very low and very high strain magnitude compared to formation sites (significant effect of adaptation type). This is likely because resorption eats into existing bone surfaces while formation builds away from existing surfaces. As the FE mesh was generated from baseline bone masks, formation sites were more likely to be distant from any given FE element.

**Fig. 7.**
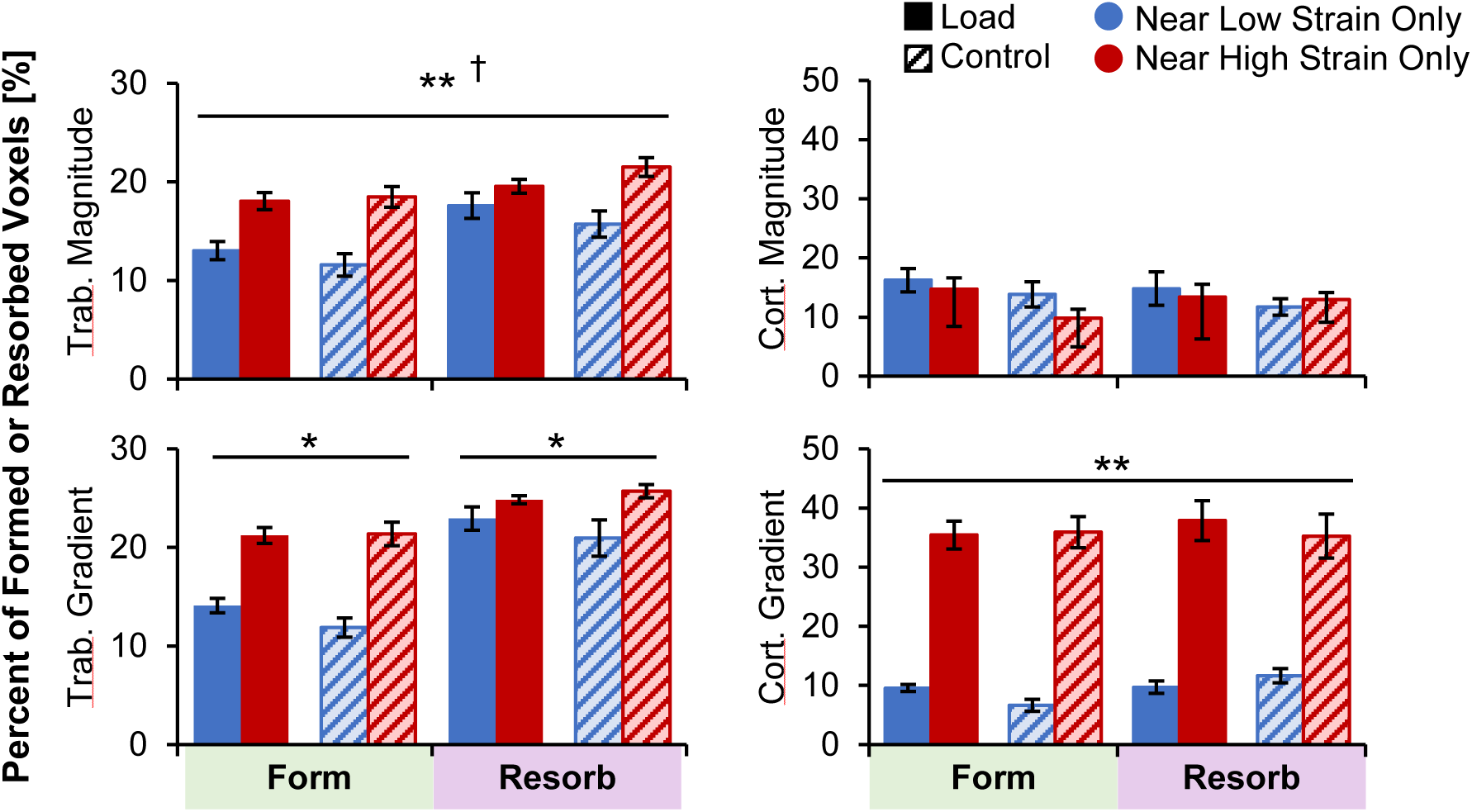
Percent of trabecular (left) and cortical (right) formation and resorption sites near very high or very low strain magnitude (top) and gradient (bottom) elements for the load (n=11) and control (n=10) groups. Data presented as group means (error bars: SEM). *Given significant strain by adaptation interaction, indicates significant difference between very low and very high strain within formation or resorption for both groups. **Indicates significant main effect of strain level (very high versus very low). †Indicates significant main effect of adaptation type (form versus resorb).

### What Percent of High or Low Strain Elements are Near Formation and Resorption Sites?

In the trabecular compartment, a greater proportion of very low strain magnitude and gradient elements were near resorption versus formation in both the load and control groups (Fig. 8). The proportion of very high strain magnitude and gradient elements near formation and resorption were similar. Therefore, trabecular bone resorption is associated with low bone strain. In the cortical compartment, very high and very low strain elements were found near formation and resorption at similar rates. Between 21-35% of low strain elements were near resorption, and between 19-43% of high strain elements were near formation. At least 12% of very low or high elements near both or neither adaptation types (Supplementary Fig. 2). There were significant interactions between group (load versus control) and adaptation type (formation versus resorption) for all metrics. For example, in the cortical compartment, both very high and low strain magnitude elements were more likely to be near formation for the load group and near resorption for the control group. These effects were driven by having more formation sites in the load group and more resorption sites in the control group, with any given element more likely to be near the more abundant adaptation type.

**Fig. 8.**
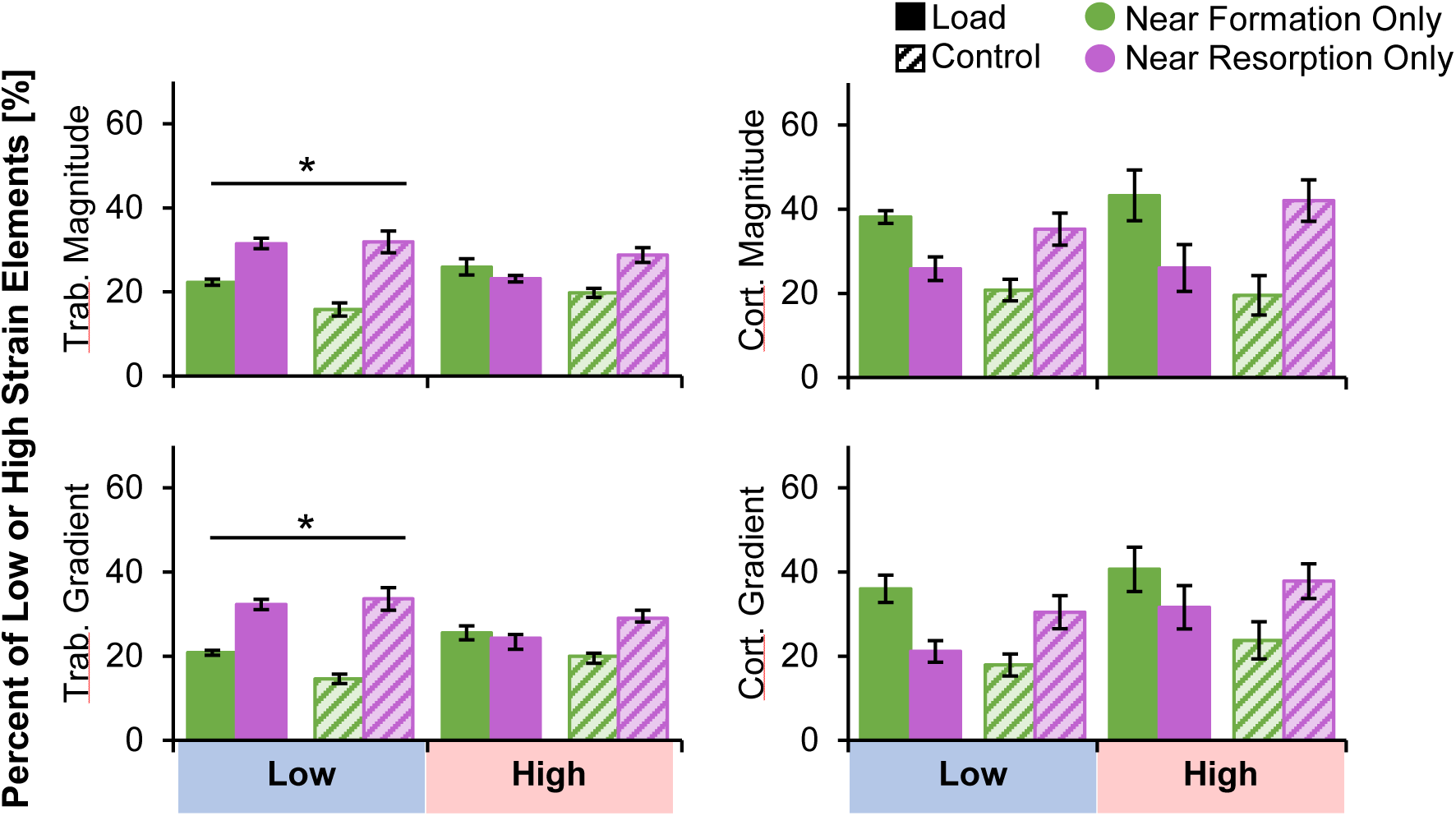
Percent of trabecular (left) and cortical (right) very low and very high strain magnitude (top) and gradient (bottom) elements near formation and resorption for the load (n=11) and control (n=10) groups. Data presented as group means (error bars: SEM). *Given significant strain by adaptation interaction, indicates significant difference between formation and resorption within very low strain elements only. Significant group by adaptation interactions were observed for all metrics but are not indicated on plot for visual clarity.

### How are Adaptation and Bone Strain Distributed around the Cortical Shell?

There were no significant differences in cortical strain magnitude or gradient between the load and control groups in any sector. Therefore, as expected, our boundary conditions based on participant-specific force recordings for the load group and average force for control subjects generated similar bone loading. The number of formation sites was higher for the load versus control group in five out of sixteen sectors (Fig. 9) located in the anterior and posterior surfaces (p<0.05). There were significantly more resorption sites in the control versus load group in one sector located in the posterior/radial quadrant.

**Fig. 9.**
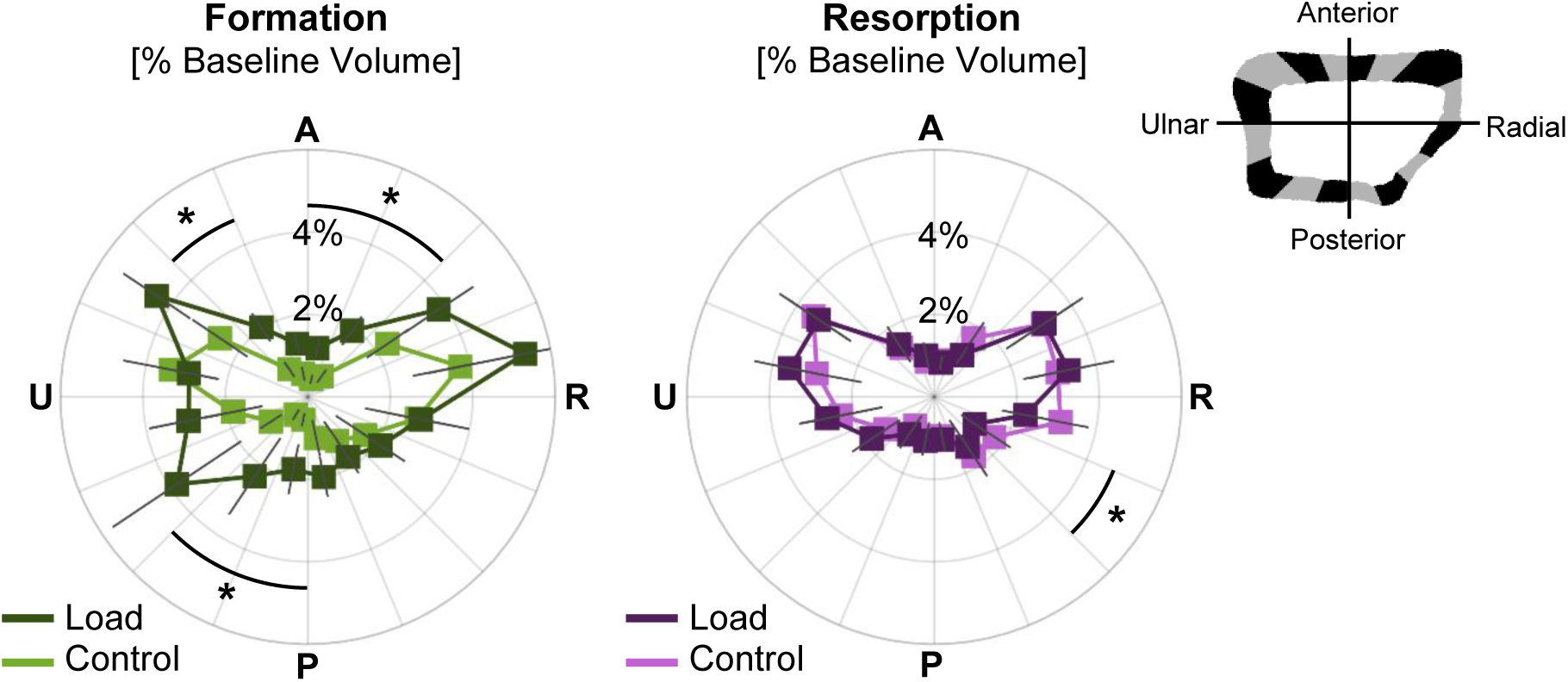
Angular distribution of formed (left) and resorbed (right) bone, as a percent of baseline cortical bone volume, in the load (n=11) and control (n=10) groups. Data presented as group means (error bars: SEM) for each sector spanning the anterior (A), ulnar (U), posterior (P), and radial (R) surfaces. *Indicates significant difference between groups.

## DISCUSSION

Our purpose was to relate tissue-level bone strain to local adaptation in the distal radius of women following 12 months of axial forearm loading. Our hypothesis that bone formation would occur preferentially in high-strain magnitude and gradient regions and bone resorption would occur preferentially in low-strain regions was partially supported. Trabecular strain magnitude and gradient were higher near formation versus resorption, and very low strain elements were more likely to be near trabecular resorption than formation. However, trabecular formation and resorption sites were both more likely to be near very high versus very low strain elements. We interpret these findings as evidence that in local regions of high strain, osteocyte stimulation and damage lead to increases in bone formation and remodeling, while in low strain regions with insufficient osteocyte stimulation, bone is removed. In the cortical compartment, the association between strain and adaptation was less clear. Strain gradient was higher near formation versus resorption for the control group, and formation and resorption were both more likely to be near very high versus low strain gradient elements. However, there were no differences in the proportion of very low and high strain elements near formation versus resorption.

Contrary to our hypothesis, similar relationships between strain and adaptation were observed in the loading and control groups. This could be interpreted to mean that, at a local level, the same mechanical cues are driving tissue remodeling, regardless of whether there was a net gain or loss in bone mass. It is unsurprising that in the absence of a novel intervention, bone adaptation is still regulated in part by bone strain generated during activities of daily living. Since the control group did not participate in the loading intervention, we did not expect measurable relationships between FE-estimated strain and adaptation because the simulated loading task was not actually performed. However, axial compression is the primary loading mode for many common activities, and the FE-estimated strain distribution may be similar to habitual strains for the control subjects. This is in agreement with Christen et al. (2014) [32], who found that in the distal tibia of postmenopausal women with normal activity levels (i.e. no intervention), formation was more frequent in regions of high strain energy density. Troy et al. (2018) found that FE-estimated principal stresses predicted four-year circum-menarcheal changes in total BMC and cortical thickness in the distal radius of non-gymnasts, but not gymnasts [42], further suggesting that bone adaptation is related to loading even when activity levels are not above those of daily living. Additionally, by defining low and high strain elements based on 5^th^ and 95^th^ percent values within an individual participant, we did not define absolute strain “setpoints” across subjects, which may vary between individuals based on activity level and other physiological factors.

Our previous work quantified the relationship between bone strain and adaptation at the macrostructural level in a pilot group of 23 women who completed fourteen weeks of forearm loading [26]. Strain was estimated by continuum-only FE models and changes in bone volume, density, and mineral content were measured from clinical resolution CT scans. When a 3 cm transverse section of the distal radius was divided into 12 subregions, there was a significant correlation between strain and change in density (but not volume or mineral content) for the load group only. In the present study, we found significant associations between strain and adaptation in the trabecular compartment, but did not detect many differences in the relationship of strain versus adaptation between the load and control groups. Continuum strains and micro-FE derived strains cannot be directly compared, and the local regions of formation/resorption measured in the present study cannot be compared to regional averages combining the trabecular and cortical compartments. Additionally, the analyses differed in duration (14 weeks versus 12 months), and the present analysis was limited to the overlapping region between baseline and follow-up HRpQCT scans (maximum 9.02 mm transverse region), while our previous analysis covered a larger, 3 cm region.

Our findings for the trabecular compartment are generally consistent with previous work in animal models, but the strength of observed differences were smaller. In the mouse caudal vertebral loading model, Schulte et al. (2013) [43] found a 39% difference in strain energy density (SED) in regions with formation versus resorption after four weeks of loading, and Lambers et al. (2015) [44] found over a 100% difference after six weeks. In comparison, we found that median trabecular energy equivalent strain magnitude and gradient were 5-8% higher in regions with formation versus resorption. In the present analysis, because all elements in micro-FE portion of our models had the same modulus and size, SED and energy equivalent strain are directly related, with SED proportional to the square of energy equivalent strain. Therefore, the direction of differences in SED and energy equivalent strains near formation and resorption should be similar, with differences in SED likely magnified compared to energy equivalent strain because of their mathematical definitions. Looking at remodeling probabilities, Cresswell et al. (2016) [21] found that after one week of vertebral loading in mice, 47% of high SED regions (defined as top 20%) were within 25 microns of bone formation. After four weeks, Schulte et al. (2013) found conditional probabilities of formation and resorption at high and low SED regions, respectively, were between 40-50%. In our participants, at most 33% of very low trabecular strain elements were near resorption only, and 23% of very high trabecular strain elements were near formation only. One explanation for the weaker relationships between strain and adaptation in our study is that adaptation in animal models is measured using micro-CT, which has a higher resolution than HRpQCT with a typical voxel size of 10-25 µm. Therefore, the amount of erroneously labeled adaptation due to partial volume effect is likely higher using HRpQCT (82 µm voxel size), limiting the strength of the measurable relationship.. Additionally, there are several physiological and lifestyle factors that cannot be controlled but likely influence bone adaptation in humans. Further work is needed to determine the individual roles that age, physical activity, calcium and Vitamin D intake, genetics, and hormonal factors plan in adaptation of bone to mechanical loading.

The current analysis did not find consistent evidence that cortical bone changes were directly related to bone strain magnitude. The 75^th^ percentile of cortical strain gradient was significantly higher for formation versus resorption, but the proportion of very low and high strain elements near formation and resorption sites were similar. We found that significantly more formation than resorption sites were located near high cortical strain gradients in both the control and loading groups. This is likely driven by the fact that cortical strain gradients are largest at the periosteal surface due to the presence of the surface itself, and that bone formation occurs on bone surfaces. Taken together, these findings suggest that cortical strain gradient, rather than magnitude, is related to adaptation. This is in agreement with loading studies in a turkey ulna exogenous loading model [17] and rooster tarsometatarsus running model [16]. In these models, circumferential strain gradients, but not strain magnitude, predicted areas of new bone formation with R^2^ values between 0.36 and 0.63. The lack of definitive strain-adaptation relationships observed in the cortical compartment in the current study may also be related to the intervention being overall more osteogenic in the trabecular compartment. The macrostructural changes, described in detail elsewhere [29], showed that the largest loading related changes were in trabecular density, especially in the inner 60% of the trabecular compartment. Additionally, we found that across subjects, age was negatively correlated with trabecular density at baseline [30], even within our relatively young age range of 21-40. Therefore, it is possible that in this group, trabecular bone is the first to undergo age-related deterioration and has the most potential for improvement due to mechanical loading. Furthermore, trabecular bone has abundant surface area available for bone apposition, in contrast to the relatively limited cortical shell. Finally, cortical bone may be at its physiological maximum, or changes in the cortical compartment may be dominated by physiological factors not directly controlled here that overshadow the influence of loading.

This study has several limitations. First, the labeling of adaptation from HRpQCT images is subject partial volume and registration error, which leads to some bone being erroneously labeled as formation or resorption. Our 10-11% error rates in the trabecular compartment are approximately double those reported by Schulte et al. using micro-CT with a 10.5 µm voxel size [43]. While micro-CT is not safe for use in humans, future studies may be improved by using newer, higher resolution HRpQCT scanners, which currently scan at a 61 µm voxel size [45]. Looking at other technologies used in humans, clinical resolution CT can be registered to measure regional changes in apparent density and bone mineral content with high repeatability (coefficient of variation <0.7% in [26]), but cannot capture adaptation at the sub-millimeter, tissue level. Despite relatively high short-term precision errors, we found that the amount of formation and resorption observed longitudinally was at least 1.7 times that for a short-term precision data set. Additionally, there was no bias toward mislabeling formation or resorption, suggesting that precision error may limit the strength, but not the direction of the measurable relationship between strain and adaptation. Overall, this supports the validity of the significant trends we have observed, but it is possible that we have underestimated absolute differences in strain parameters and the spatial association between strain and adaptation. Second, our findings may overestimate strain/adaptation relationships due to selection of “responders” within the loading group. However, the relationships between local strain and adaptation type were consistent in both loading and control subjects. Third, the FE model boundary conditions used participant-specific load magnitudes but assumed an axial direction. While we instruct participants to perform loading with their arm directed axially, measuring the exact positioning was outside the scope of this investigation. The potential influence of variability in loading position on FE-estimated bone strain is part of our ongoing work. We focused on the magnitude and norm of the spatial gradient of energy equivalent strain, as energy equivalent strain is a scalar representative of the multiaxial strain state, and has been shown to relate to adaptation in our pilot study [24]. Additionally, the norm of strain gradients in the axial and transverse directions as a scalar representation of spatial variability has been used in previous studies [39,46]. However, it is possible that other bone strain parameters have a controlling role in functional bone adaptation. Bone tissue strain gradient was selected as an upstream measure of fluid flow, as spatially varying strains yield pressure gradients and therefore flow within lacunar-canicular and marrow spaces. A more direct estimate of fluid flow using poroelastic modeling [47–49] or inclusion of marrow as a separate material [19] may provide a more detailed description of the local mechanical environment and have shown potential in predicting adaptation in animal loading models, but is outside the scope of the present study.

In summary, we related tissue-level bone strain to 12-month changes in radius microstructure in young healthy women who performed axial forearm loading or participated as non-loading controls. We found that local regions of high strain magnitude and gradient are associated with increased trabecular formation and remodeling, while low strain magnitude and gradient are associated with trabecular bone resorption. Cortical strain gradient was higher near formation versus resorption in the control group, and both adaptation types occurred preferentially near high strain gradients at the periosteal surface. While we observed a significant measurable relationship between strain and adaptation, only half of very high and low strain elements were near formation or resorption only. The similarity of the strain/adaptation relationship between loading and control groups suggest that, at a local level, the same mechanical cues drive tissue remodeling, regardless of the net stimulus or change. Overall, our results show that participant-specific bone strain and adaptation can be estimated using currently available non-invasive imaging techniques. Our results also highlight that bone strain has a measurable, controlling influence on the adaptive response in healthy adult women. To the best of our knowledge, this is the first study to relate prospectively measured changes in human bone structure to subject-specific bone strain, based on real force measurements. This is an important first step toward defining loading thresholds above or below which bone formation or resorption occur, quantifying the extent to which changes in human bone structure can be predicted based on strain, and characterizing the influence of physical activity history, age, and other physiological factors on these thresholds. Ultimately, such knowledge could inform predictive models of bone adaptation, enabling the *in silico* comparison and optimization of targeted loading interventions to maximize bone strength and prevent fragility fractures.

## ACKNOWLEDGMENT

We thank Joshua Johnson, Tifiny Butler, and Sabahat Ahmed for their role in data collection for the parent study. We thank Michael DiStefano for his assistance with local adaptation image analysis.

## FUNDING

This work was supported by NIAMS of the National Institutes of Health under award number R01AR063691. This material is based upon work supported by the National Science Foundation Graduate Research Fellowship Program under Grant No. DGE-1106756. This research was performed using computational resources supported by the Academic & Research Computing group at Worcester Polytechnic Institute and the National Science Foundation under Grant No. DMS-1337943. Any opinions, findings, and conclusions or recommendations expressed in this material are those of the author(s) and do not necessarily reflect the views of the National Science Foundation or the National Institutes of Health.

## Supplementary Figure Captions List

**Fig. S1.**
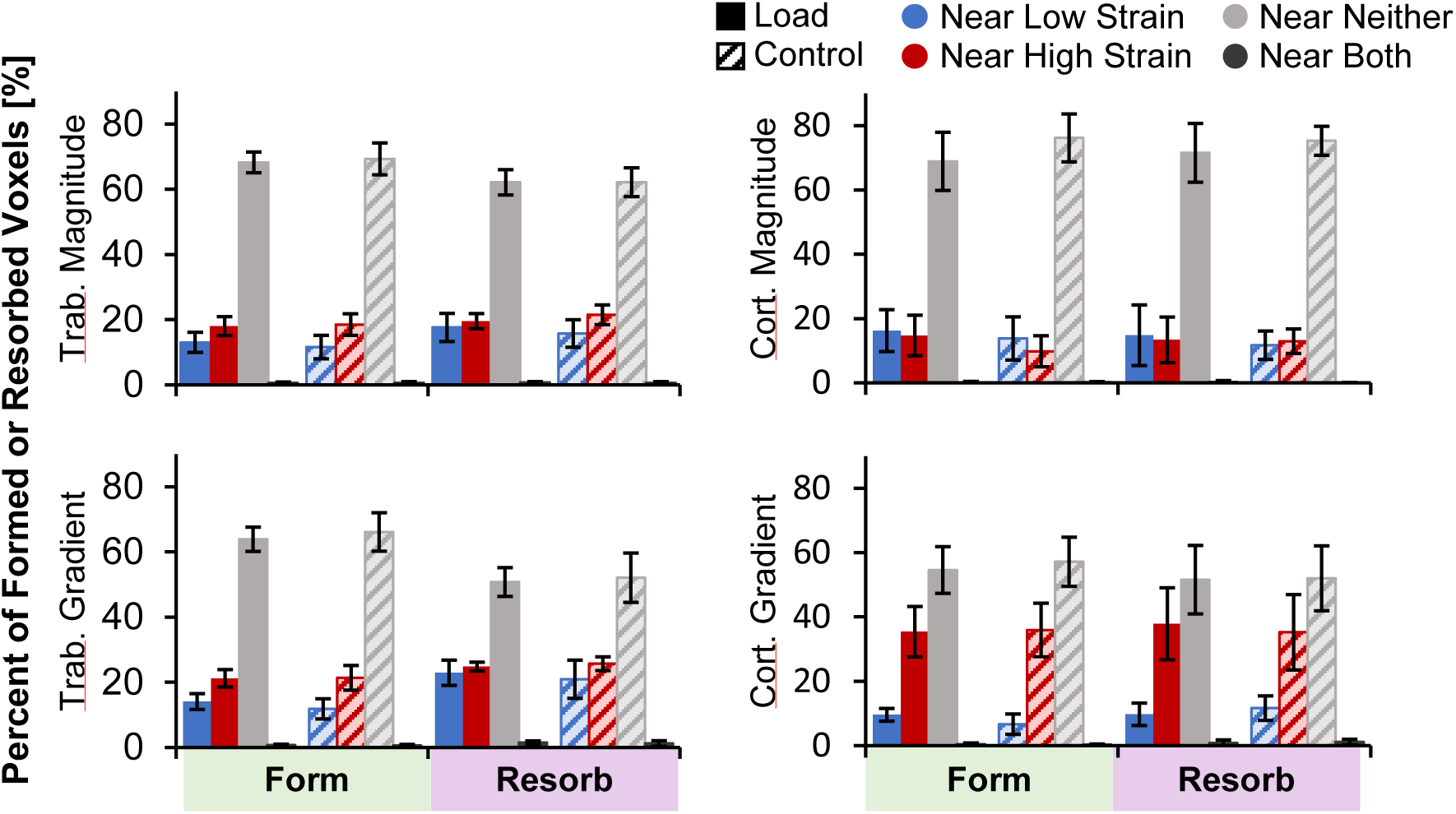
Percent of trabecular (left) and cortical (right) formation and resorption sites near very high only, very low only, neither very high nor very low, or both very high and very low strain magnitude (top) and gradient (bottom) elements for the load (n=11) and control (n=10) groups. Data presented as group means (error bars: SEM).

**Fig. S2.**
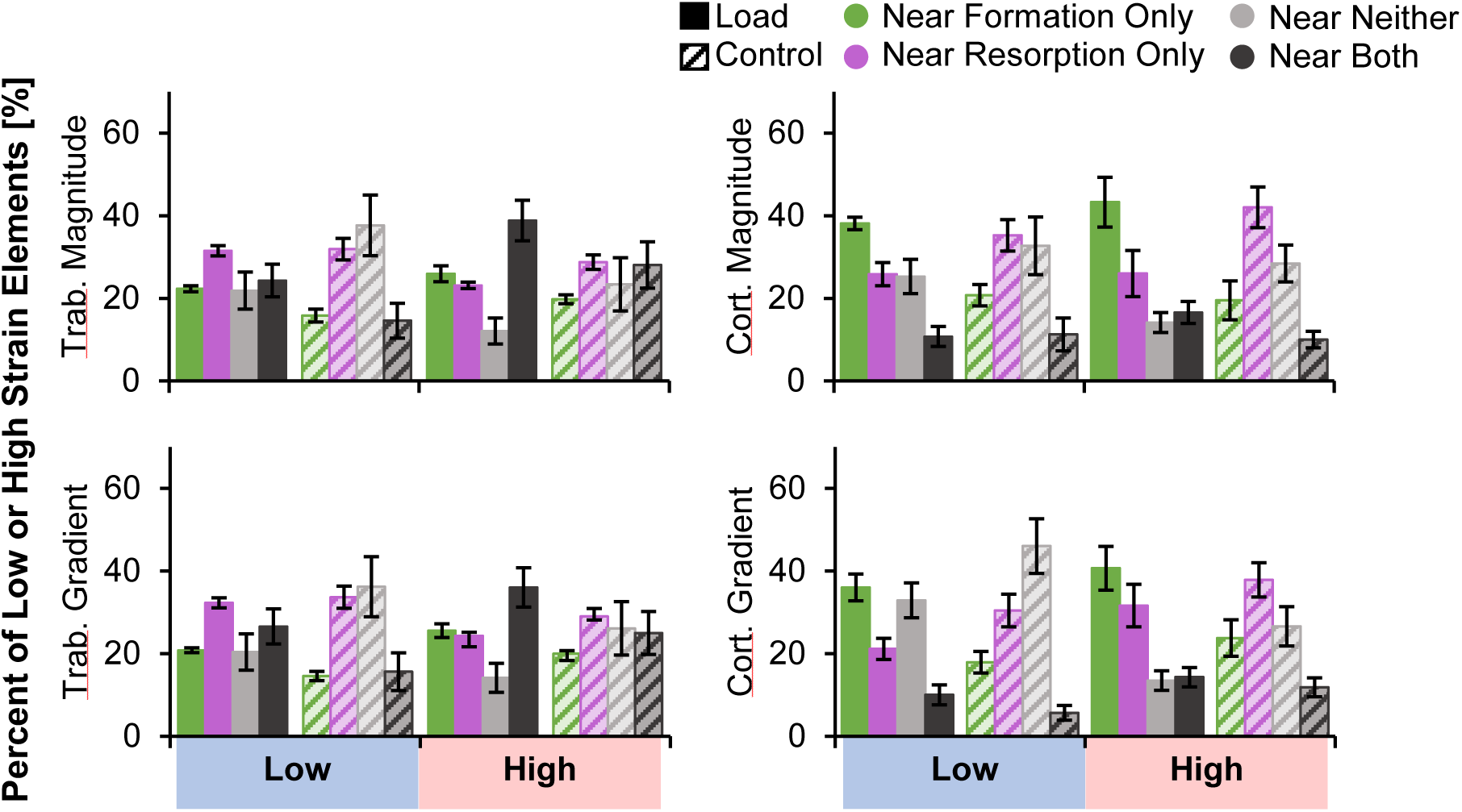
Percent of trabecular (left) and cortical (right) very low and very high strain magnitude (top) and gradient (bottom) elements near formation only, near resorption only, near neither formation nor resorption, or near both formation and resorption for the load (n=11) and control (n=10) groups. Data presented as group means (error bars: SEM).

## Supplementary Table Caption List

**Table S1:** Mean (SD) strain magnitude and strain gradient near formation and resorption sites, in the trabecular and cortical compartments, for the load and control groups. Differences reflect the mean (SD) values for within-participant differences in strain near formation and resorption, with positive differences indicating higher strain near formation.

|  |  |  | Form | Resorb | Difference |
| --- | --- | --- | --- | --- | --- |
| Trabecular | Magnitude ($\mu\epsilon$) | Load | 196.6 (60.9) | 187.7 (59.5) | 9.0 (12.8) |
|  |  | Control | 232.4 (36.4) | 217.5 (26.2) | 14.9 (21.3) |
| | Gradient ($\mu\epsilon/\text{mm}$) | Load | 587.9 (187.2) | 548.8 (174.3) | 39.1 (34.2) |
|  |  | Control | 669.4 (100.0) | 616.3 (107.9) | 53.1 (43.5) |
| Cortical | Magnitude ($\mu\epsilon$) | Load | 280.7 (90.6) | 297.5 (87.6) | -16.7 (49.9) |
|  |  | Control | 299.5 (77.6) | 314.7 (77.4) | -15.2 (36.4) |
| | Gradient ($\mu\epsilon/\text{mm}$) | Load | 353.6 (157.8) | 317.4 (117.6) | 36.2 (101.5) |
|  |  | Control | 443.3 (146.6) | 304.4 (86.7) | 138.9 (98.6) |

## Notes

### Competing Interest Statement

The authors have declared no competing interest.

## REFERENCES

[1] 2017, “National Osteoporosis Foundation. Osteoporosis Exercise for Strong Bones.,” pp. 1–3 [Online]. Available: https://www.nof.org/patients/fracturesfall-prevention/exercisesafe-movement/osteoporosis-exercise-for-strong-bones/. [Accessed: 03-Mar-2018].

[2] Bareither, M. Lou, Grabiner, M. D., and Troy, K. L., 2008, “Habitual Site-Specific Upper Extremity Loading Is Associated with Increased Bone Mineral of the Ultradistal Radius in Young Women,” J. Women’s Heal., 17(10), pp. 1577–1581.

[3] Stewart, A. A. D., and Hannan, J., 2000, “Total and Regional Bone Density in Male Runners, Cyclists, and Controls,” Med Sci Sport. Exerc, 32(17), pp. 1373–1377.

[4] Kontulainen, S., Sievänen, H., Kannus, P., Pasanen, M., and Vuori, I., 2003, “Effect of Long-Term Impact-Loading on Mass, Size, and Estimated Strength of Humerus and Radius of Female Racquet-Sports Players: A Peripheral Quantitative Computed Tomography Study between Young and Old Starters and Controls,” J. Bone Miner. Res., 18(2), pp. 352–359.

[5] Bass, S. L., Saxon, L., Daly, R. M., Turner, C. H., Robling, A. G., Seeman, E., and Stuckey, S., 2002, “The Effect of Mechanical Loading on the Size and Shape of Bone in Pre-, Peri-, and Postpubertal Girls: A Study in Tennis Players,” J. Bone Miner. Res., 17(12), pp. 2274–2280.

[6] Warden, S. J., Carballido-gamio, J., Avin, K. G., Kersh, M. E., Fuchs, R. K., Krug, R., and Bice, R. J., 2019, “Adaptation of the Proximal Humerus to Physical Activity : A within-Subject Controlled Study in Baseball Players,” Bone, 121(October 2018), pp. 107–115.

[7] Zhao, R., Zhang, M., and Zhang, Q., 2017, “The Effectiveness of Combined Exercise Interventions for Preventing Postmenopausal Bone Loss: A Systematic Review and Meta-Analysis,” J. Orthop. Sport. Phys. Ther., 47(4), pp. 241–251.

[8] Zhao, R., Zhao, M., and Zhang, L., 2014, “Efficiency of Jumping Exercise in Improving Bone Mineral Density Among Premenopausal Women: A Meta-Analysis,” Sport. Med., 44(10), pp. 1393–1402.

[9] Martyn St James, M., and Carroll, S., 2010, “Effects of Different Impact Exercise Modalities on Bone Mineral Density in Premenopausal Women: A Meta-Analysis,” J. Bone Miner. Metab., 28(3), pp. 251–267.

[10] Wallace, B. A., and Cumming, R. G., 2000, “Systematic Review of Randomized Trials of the Effect of Exercise on Bone Mass in Pre- and Postmenopausal Women.,” Calcif. Tissue Int., 67(1), pp. 10–8.

[11] Goodship, A. E., Lanyon, L. E., and McFie, H., 1979, “Functional Adaptation of Bone to Increased Stress. An Experimental Study,” J. Bone Jt. Surg. - Ser. A.

[12] Rubin, C. T., and Lanyon, L. E., 1985, “Regulation of Bone Mass by Mechanical Strain Magnitude,” Calcif. Tissue Int., 37(4), pp. 411–417.

[13] Rubin, C. T., and Lanyon, L. E., 1984, “Regulation of Bone Formation by Applied Dynamic Loads,” J. Bone Jt. Surg., 66A(3), pp. 397–402.

[14] Lanyon, L. E., Goodship, A. E., Pye, C. J., and MacFie, J. H., 1982, “Mechanically Adaptive Bone Remodelling,” J. Biomech., 15(3), pp. 141–154.

[15] Kotha, S. P., Hsieh, Y. F., Strigel, R. M., Müller, R., and Silva, M. J., 2004, “Experimental and Finite Element Analysis of the Rat Ulnar Loading Model - Correlations between Strain and Bone Formation Following Fatigue Loading,” J. Biomech., 37(4), pp. 541–548.

[16] Judex, S., Gross, T. S., and Zernicke, R. F., 1997, “Strain Gradients Correlate with Sites of Exercise-Induced Bone-Forming Surfaces in the Adult Skeleton,” J. Bone Miner. Res., 12(10), pp. 1737–1745.

[17] Gross, T. S., Edwards, J. L., Mcleod, K. J., and Rubin, C. T., 1997, “Strain Gradients Correlate with Sites of Periosteal Bone Formation,” J. Bone Miner. Res., 12(6), pp. 982–988.

[18] Kim, C. H., Takai, E., Zhou, H., Von Stechow, D., Muller, R., Dempster, D. W., and Guo, X. E., 2003, “Trabecular Bone Response to Mechanical and Parathyroid Hormone Stimulation: The Role of Mechanical Microenvironment,” J. Bone Miner. Res., 18(12), pp. 2116–2125.

[19] Webster, D., Schulte, F. A., Lambers, F. M., Kuhn, G., and Müller, R., 2015, “Strain Energy Density Gradients in Bone Marrow Predict Osteoblast and Osteoclast Activity: A Finite Element Study,” J. Biomech., 48(5), pp. 866–874.

[20] Schulte, F. A., Zwahlen, A., Lambers, F. M., Kuhn, G., Ruffoni, D., Betts, D., Webster, D. J., and M??ller, R., 2013, “Strain-Adaptive in Silico Modeling of Bone Adaptation - A Computer Simulation Validated by in Vivo Micro-Computed Tomography Data,” Bone, 52(1), pp. 485–492.

[21] Cresswell, E. N., Goff, M. G., Nguyen, T. M., Lee, W. X., and Hernandez, C. J., 2016, “Spatial Relationships between Bone Formation and Mechanical Stress within Cancellous Bone,” J. Biomech., 49(2), pp. 222–228.

[22] Paul, G. R., Malhotra, A., and Müller, R., 2018, “Mechanical Stimuli in the Local In Vivo Environment in Bone: Computational Approaches Linking Organ-Scale Loads to Cellular Signals,” Curr. Osteoporos. Rep., 16(4), pp. 395–403.

[23] Hinton, P. V, Rackard, S. M., Kennedy, O. D., and Kennedy, O. D., 2018, “In Vivo Osteocyte Mechanotransduction : Recent Developments and Future Directions,” pp. 746–753.

[24] Troy, K. L., Edwards, W. B., Bhatia, V. A., and Bareither, M. Lou, 2013, “In Vivo Loading Model to Examine Bone Adaptation in Humans: A Pilot Study,” J. Orthop. Res., 31(9), pp. 1406–1413.

[25] Bhatia, V. A., Edwards, W. B., and Troy, K. L., 2014, “Predicting Surface Strains at the Human Distal Radius during an in Vivo Loading Task - Finite Element Model Validation and Application,” J. Biomech., 47(11), pp. 2759–2765.

[26] Bhatia, V. A., Brent Edwards, W., Johnson, J. E., and Troy, K. L., 2015, “Short-Term Bone Formation Is Greatest Within High Strain Regions of the Human Distal Radius: A Prospective Pilot Study,” J. Biomech. Eng., 137(1), pp. 011001-1–5.

[27] Laib, A., Hauselmann, H. J., and Ruegsegger, P., 1998, “In Vivo High Resolution 3D-QCT of the Human Forearm,” Technol. Heal. Care, 6, pp. 329–337.

[28] Johnson, J. E., and Troy, K. L., 2017, “Validation of a New Multiscale Finite Element Analysis Approach at the Distal Radius,” Med. Eng. Phys., 44, pp. 16–24.

[29] Troy, K. L., Mancuso, M. E., Johnson, J. E., Wu, Z., Schnitzer, T. J., and Butler, T. A., 2020, “Bone Adaptation in Adult Women Is Related to Loading Dose: A 12-month Randomized Controlled Trial.,” J. Bone Miner. Res., p. jbmr.3999.

[30] Mancuso, M. E., Johnson, J. E., Ahmed, S. S., Butler, T. A., and Troy, K. L., 2018, “Distal Radius Microstructure and Finite Element Bone Strain Are Related to Site-Specific Mechanical Loading and Areal Bone Mineral Density in Premenopausal Women,” Bone Reports, 8(July 2017), pp. 187–194.

[31] Pialat, J. B., Burghardt, A. J., Sode, M., Link, T. M., and Majumdar, S., 2012, “Visual Grading of Motion Induced Image Degradation in High Resolution Peripheral Computed Tomography: Impact of Image Quality on Measures of Bone Density and Micro-Architecture,” Bone, 50(1), pp. 111–118.

[32] Christen, P., Ito, K., Ellouz, R., Boutroy, S., Sornay-Rendu, E., Chapurlat, R. D., and van Rietbergen, B., 2014, “Bone Remodelling in Humans Is Load-Driven but Not Lazy,” Nat. Commun., 5, p. 4855.

[33] Christen, P., Boutroy, S., Ellouz, R., Chapurlat, R., and Van Rietbergen, B., 2018, “Least-Detectable and Age-Related Local in Vivo Bone Remodelling Assessed by Time-Lapse HR-PQCT,” PLoS One, 13(1), pp. 1–11.

[34] MacNeil, J. A., and Boyd, S. K., 2007, “Accuracy of High-Resolution Peripheral Quantitative Computed Tomography for Measurement of Bone Quality.,” Med. Eng. Phys., 29(10), pp. 1096–1105.

[35] Baim, S., Wilson, C. R., Lewiecki, E. M., Luckey, M. M., Downs, R. W., and Lentle, B. C., 2005, “Precision Assessment and Radiation Safety for Dual-Energy X-Ray Absorptiometry,” J. Clin. Densitom.

[36] Morgan, E. F., Bayraktar, H. H., and Keaveny, T. M., 2003, “Trabecular Bone Modulus-Density Relationships Depend on Anatomic Site,” J. Biomech., 36(7), pp. 897–904.

[37] Anderson, D. D., Deshpande, B. R., Daniel, T. E., and Baratz, M. E., 2005, “A Three-Dimensional Finite Element Model of the Radiocarpal Joint: Distal Radius Fracture Step-off and Stress Transfer.,” Iowa Orthop. J., 25, pp. 108–117.

[38] Huiskes, R., Ruimerman, R., van Lenthe, G. H., and Janssen, J. D., 2000, “Effects of Mechanical Forces on Maintenance and Adaptation of Form in Trabecular Bone.,” Nature, 405(6787), pp. 704–706.

[39] Ruimerman, R., Van Rietbergen, B., Hilbers, P., and Huiskes, R., 2005, “The Effects of Trabecular-Bone Loading Variables on the Surface Signaling Potential for Bone Remodeling and Adaptation,” Annals of Biomedical Engineering, pp. 71–78.

[40] Adachi, T., Tomita, Y., Sakuae, H., and Tanaka, M., 1997, “Simulation of Trabecular Surface Remodeling Based on Local Stress Nonuniformity,” JSME Int. J. Ser. C Mech. Syst. Mach. Elem. Manuf., 40(4), pp. 415–434.

[41] Morgan, T. G., Bostrom, M. P. G., and van der Meulen, M. C. H., 2015, “Tissue-Level Remodeling Simulations of Cancellous Bone Capture Effects of in Vivo Loading in a Rabbit Model,” J. Biomech., 48(5), pp. 875–882.

[42] Troy, K. L., Scerpella, T. A., and Dowthwaite, J. N., 2018, “Circum-Menarcheal Bone Acquisition Is Stress-Driven: A Longitudinal Study in Adolescent Female Gymnasts and Non-Gymnasts,” J. Biomech., 78, pp. 45–51.

[43] Schulte, F. A., Ruffoni, D., Lambers, F. M., Christen, D., Webster, D. J., Kuhn, G., and Müller, R., 2013, “Local Mechanical Stimuli Regulate Bone Formation and Resorption in Mice at the Tissue Level,” PLoS One, 8(4).

[44] Lambers, F. M., Kuhn, G., Weigt, C., Koch, K. M., Schulte, F. A., and Müller, R., 2015, “Bone Adaptation to Cyclic Loading in Murine Caudal Vertebrae Is Maintained with Age and Directly Correlated to the Local Micromechanical Environment,” J. Biomech., 48(6), pp. 1179–1187.

[45] Manske, S. L., Zhu, Y., Sandino, C., and Boyd, S. K., 2015, “Human Trabecular Bone Microarchitecture Can Be Assessed Independently of Density with Second Generation HR-PQCT,” Bone, 79, pp. 213–221.

[46] Koontz, J. T., Charras, G. T., and Guldberg, R. E., 2002, “A Microstructural Finite Element Simulation of Mechanically Induced Bone Formation,” J. Biomech. Eng., 123(6), p. 607.

[47] Scheiner, S., Pivonka, P., and Hellmich, C., 2016, “Poromicromechanics Reveals That Physiological Bone Strains Induce Osteocyte-Stimulating Lacunar Pressure,” Biomech. Model. Mechanobiol., 15(1), pp. 9–28.

[48] Pereira, A. F., Javaheri, B., Pitsillides, A. A., and Shefelbine, S. J., 2015, “Predicting Cortical Bone Adaptation to Axial Loading in the Mouse Tibia,” J. R. Soc. Interface, 12(110).

[49] Carriero, A., Pereira, A. F., Wilson, A. J., Castagno, S., Javaheri, B., Pitsillides, A. A., Marenzana, M., and Shefelbine, S. J., 2018, “Spatial Relationship between Bone Formation and Mechanical Stimulus within Cortical Bone: Combining 3D Fluorochrome Mapping and Poroelastic Finite Element Modelling,” Bone Reports, 8(February), pp. 72–80.

